# Shared functional connections within and between cortical networks predict cognitive abilities in adult males and females

**DOI:** 10.1101/2021.02.17.431670

**Authors:** Elvisha Dhamala, Keith W. Jamison, Abhishek Jaywant, Amy Kuceyeski

## Abstract

A thorough understanding of sex-independent and sex-specific neurobiological features that underlie cognitive abilities in healthy individuals is essential for the study of neurological illnesses in which males and females differentially experience and exhibit cognitive impairment. Here, we evaluate sex-independent and sex-specific relationships between functional connectivity and individual cognitive abilities in 392 healthy young adults (196 males) from the Human Connectome Project. First, we establish that sex-independent models comparably predict crystallised abilities in males and females, but more accurately predict fluid abilities in males. Second, we demonstrate sex-specific models comparably predict crystallised abilities within and between sexes, and generally fail to predict fluid abilities in either sex. Third, we reveal that largely overlapping connections between visual, dorsal attention, ventral attention, and temporal parietal networks are associated with better performance on crystallised and fluid cognitive tests in males and females, while connections within visual, somatomotor, and temporal parietal networks are associated with poorer performance. Together, our findings suggest that shared neurobiological features of the functional connectome underlie crystallised and fluid abilities across the sexes.

## Introduction

Sex differences in brain-behaviour relationships are widely studied and controversial in neuroscience. Studies often report contradictory findings, and many are not replicated. While some studies have found evidence for sex differences in healthy cognitive functioning (Camarata & Woodcock, 2006; Irwing & Lynn, 2005; Lynn & Irwing, 2004), others have reported a lack of differences (Hyde, 2005; Jäncke, 2018). Similarly, while some studies have found evidence for sex differences in healthy brain function and structure (Cummings et al., 2020; De Bellis et al., 2001; Ruben C Gur et al., 1999; Ingalhalikar et al., 2014; Kogler et al., 2016; Ritchie et al., 2018; Rodriguez, Warkentin, Risberg, & Rosadini, 1988; Scheinost et al., 2015; Weis, Hodgetts, & Hausmann, 2019; Weis, Patil, et al., 2019), others have observed the opposite (Bishop & Wahlsten, 1997; Eliot, Ahmed, Khan, & Patel, 2021; Sommer, Aleman, Somers, Boks, & Kahn, 2008). Finally, while some studies have demonstrated sex differences in the relationship between neural function/structure and cognitive functioning in healthy individuals (R. C. Gur & Gur, 2017; Kimura, 2004; Satterthwaite et al., 2015), others have shown otherwise (Eliot, 2011; Sommer et al., 2008). It has also been suggested that sex differences in neural circuitry and/or neurochemistry may reflect compensation for genetic and/or hormonal differences to ensure that male and female behaviours are more similar than different , and many of the contradictory findings may be attributable to differences in sample sizes, methodology, and publication bias (Eliot et al., 2021). Hence, it remains to be determined whether males and females have shared or distinct brain-behaviour relationships.

In recent years, sex differences in cognitive manifestations of various neurological, neurodevelopmental, and neuropsychiatric illnesses have become increasingly evident (Han et al., 2012; Irvine, Laws, Gale, & Kondel, 2012; Laws, Irvine, & Gale, 2016; Subramaniapillai, Almey, Rajah, & Einstein, 2020). Insight into sex-independent and sex-specific brain-behaviour relationships in healthy young adults can enable better understanding of the neurobiological underpinnings of cognitive deficits within and across sexes, paving the way for the development and implementation of personalised treatment strategies. In this study, we aim to disentangle sex-specific and sex-independent brain-behaviour relationships between resting-state functional connectivity and cognitive abilities in healthy young adults.

Resting-state functional connectivity is defined as the temporal dependence of the blood-oxygen-level dependent (BOLD) response in anatomically separate brain regions at rest (Aertsen, Gerstein, Habib, & Palm, 1989; Friston, Frith, Liddle, & Frackowiak, 1993; Martijn P Van Den Heuvel & Pol, 2010). Many studies have linked functional connectivity to cognitive functioning (Casey, Galvan, & Hare, 2005; Casey, Giedd, & Thomas, 2000; Cole, Yarkoni, Repovs, Anticevic, & Braver, 2012; Moeller, Willmes, & Klein, 2015; Park & Friston, 2013; Seeley et al., 2007; Spreng, Stevens, Chamberlain, Gilmore, & Schacter, 2010; M. P. van den Heuvel, Stam, Kahn, & Hulshoff Pol, 2009) and predicted individual cognitive abilities from functional connectivity (Chen et al., 2020; Dhamala, Jamison, Jaywant, Dennis, & Kuceyeski, 2021; He et al., 2020; J. W. Li et al., 2019; Zimmermann, Griffiths, & McIntosh, 2018). Recent work in this area has shown global signal regression, or removal of trends in the fMRI signal, improves prediction accuracy (J. W. Li et al., 2019), machine and deep learning models perform comparably (He et al., 2020), and shared network features predict scores from distinct cognitive domains (Chen et al., 2020; Dhamala et al., 2021). These studies aim to capture brain-behaviour relationships that exist between functional connectivity and cognitive abilities, but it remains unclear whether these relationships are consistent across the sexes.

Sex differences in functional connectivity have been observed across distinct populations, including North American children and adolescents with and without psychiatric illnesses, as well as North American and German healthy adults (Cummings et al., 2020; Gong, He, & Evans, 2011; R. C. Gur & Gur, 2017; Kogler et al., 2016; Satterthwaite et al., 2015; Scheinost et al., 2015; Weis, Hodgetts, et al., 2019; Zhang, Dougherty, Baum, White, & Michael, 2018). Previous work in a developmental cohort has shown males exhibit stronger inter-network connectivity, while females exhibit stronger intra-network connectivity (Satterthwaite et al., 2015). Extant literature also suggests hormonal modulation of functional connectivity (Dubol et al., 2020; Fitzgerald, Pritschet, Santander, Grafton, & Jacobs, 2020; Hjelmervik, Hausmann, Osnes, Westerhausen, & Specht, 2014; Pritschet et al., 2020; Weis, Hodgetts, et al., 2019). In terms of functional connectivity features that discriminate sex, two studies identified that connections within and between frontoparietal and default mode networks strongly contribute to the predictions (Weis, Patil, et al., 2019; Zhang et al., 2018). Together, these studies suggest sex differences exist in functional organisation of the brain, but do not address whether these differences translate into sex differences in connectivity-cognition relationships.

A recent study similar to this one investigated differences between males and females in predictability of individual intelligence quotient (IQ) and sub-domain cognitive scores using whole-brain functional connectivity (Jiang, Calhoun, Fan, et al., 2020). Their individualized prediction integrated feature selection and regression with a leave-one-out cross validation strategy, resulting in distinct functional connectivity features being selected for each interaction. They reported IQ and other cognitive scores are generally more predictable in females than they are in males, and the sex-specific models rely on distinct functional connections to make predictions. A second study from the same group used a similar approach to predict IQ in males and females using functional connectivity, cortical thickness, or both (Jiang, Calhoun, Cui, et al., 2020). The reported no differences in prediction accuracy between males and females but found that sex-specific models relied on distinct neurobiological correlates. While these findings suggest the presence of distinct brain-behaviour relationships across the sexes, their leave-one-out prediction approach, resulting in distinct features for every iteration, limits the extent to which we can compare and generalise these results because the features used are dependent on which subject is left out in the cross validation. In this current study, we aim to address this concern and expand upon this work.

Here, we study sex-independent and sex-specific brain-behaviour relationships between functional connectivity and individual cognitive abilities in 392 healthy young adults (196 male-female pairs matched for cognitive composite scores) from the Human Connectome Project (Van Essen et al., 2013). First, we quantify whether sex-independent models differ in how accurately they can predict distinct cognitive abilities from functional connectivity in males and females. Second, we quantify whether sex-specific models better predict individual cognitive abilities from functional connectivity within or between sexes. Third, we evaluate whether shared or sex-specific functional connectivity features map to cognitive abilities.

## Methods

The methods used here build upon our prior work (Dhamala et al., 2021) but the analyses presented are novel and aim to identify shared and sex-specific features that predict cognitive abilities. Our experimental workflow is shown in Figure 1. The data that support the findings of this study are openly available as part of the Human Connectome Project at https://www.humanconnectome.org/study/hcp-young-adult/document/1200-subjects-data-release (Van Essen et al., 2013). Code used to generate the results presented here are available on GitHub (https://github.com/elvisha/SexSpecificCognitivePredictions).

**Figure 1:**
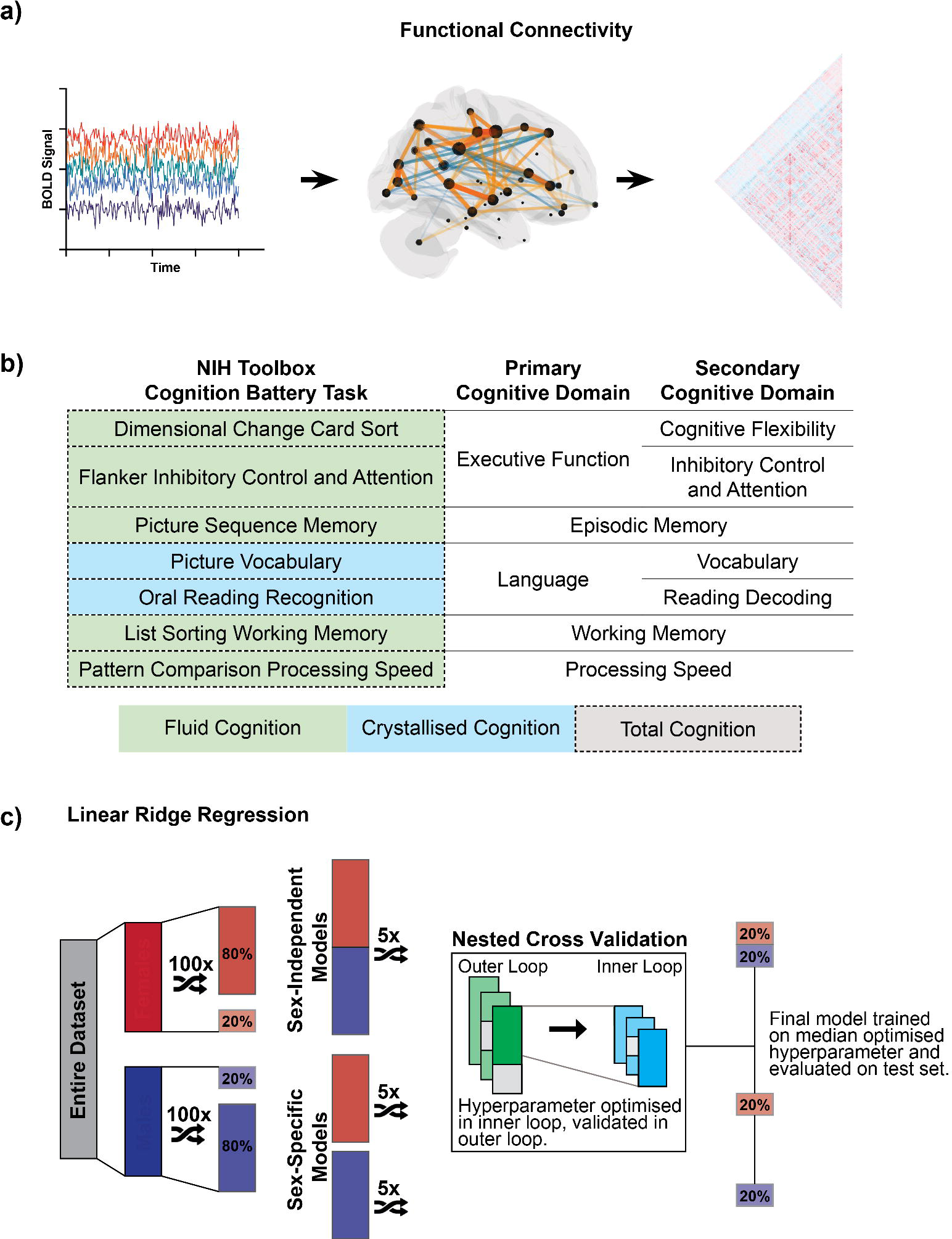
Experimental workflow. a) First, we generated individual functional connectivity using Pearson correlation of regional global signal regressed resting-state functional MRI time series. b) Second, we compiled cognitive scores for all subjects. The NIH Toolbox Cognition Battery assesses five cognitive domains using seven tests. The Crystallised Cognition Composite (blue) reflects language (vocabulary, reading decoding). The Fluid Cognition Composite (green) reflects executive function (cognitive flexibility, inhibitory control and attention), episodic memory, working memory, and processing speed. The Total Cognition Composite (dotted) combines the Crystallised and Fluid Composite scores. c) Third, we predicted each cognitive score from functional connectivity using sex-independent and sex-specific linear ridge regression models. We randomly shuffled and split the male and female subjects into train (80%) and test (20%) groups. Male and female training subsets were concatenated for the sex-independent models and kept separate for the sex-specific models. We performed five shuffled iterations of nested cross validation with three-fold inner and outer loops. The model hyperparameter was optimised in the inner loop and validated in the outer loop. The median optimised hyperparameter from five iterations of nested cross validation was used to train the final model on the entire (sex-independent or sex-specific) training set and evaluated on the (sex-independent or sex-specific) test hold-out set. This was repeated for 100 unique train/test splits.

### Dataset

We used publicly-available high resolution, preprocessed MRI data from the Human Connectome Project (HCP) – Young Adult S1200 release (Van Essen et al., 2013). MRI data were acquired on a Siemens Skyra 3T scanner at Washington University in St. Louis. Acquisitions included T1-weighted and T2-weighted anatomical images (0.7mm isotropic), and functional MRI (2.0mm isotropic, TR/TE = 720/33.1ms, 8x multiband acceleration). Functional MRI were collected with both left-right and right-left phase encoding. We examined resting-state functional MRI (rfMRI) time series from 196 male-female pairs (n=392) of unrelated healthy young adults with four complete rfMRI runs. Male-female pairs were matched for their Crystallised, Fluid, and Total composite scores to ensure there were no significant differences in cognitive function (p>0.05) between the two sexes. Although the term *gender* is used in the HCP Data Dictionary, we use the term *sex* in this article because the database collected self-reported biological sex information as opposed to gender identification. We did not verify the self-reported biological sex using genetic information.

### Parcellation

We used a subject-specific CoCo439 parcellation that was developed inhouse by combining parts of several atlases. This parcellation includes 358 (of 360) functionally derived cortical regions from HCP multi-modal parcellation (MMP) (Matthew F Glasser et al., 2016) (two hippocampal regions were excluded as they were included in other subcortical ROIs); 12 anatomically defined subcortical regions derived from FreeSurfer’s aseg.mgz, adjusted by FSL’s FIRST tool (Patenaude, Smith, Kennedy, & Jenkinson, 2011); 12 anatomically defined subcortical nuclei from AAL3v1 (Rolls, Huang, Lin, Feng, & Joliot, 2020); 30 anatomically defined subcortical nuclei from FreeSurfer 7 (Iglesias et al., 2018) (50 nuclei were merged down to 30 to remove the smallest nuclei, as with AAL3v1); and 27 anatomically defined cerebellar regions from the SUIT atlas (Diedrichsen, Balsters, Flavell, Cussans, & Ramnani, 2009). Additional details and corresponding files for this parcellation are available on GitHub (https://github.com/kjamison/nemo#parcellations).

### Functional Connectivity Extraction

Each subject underwent four gradient-echo EPI resting-state fMRI (rsfMRI) runs of ∼15 min each over two sessions. There are 1200 volumes per scan for a total of 4800 volumes for each subject over the four runs. The minimal preprocessing pipeline performed by the HCP consortium included motion and distortion correction, registration to subject anatomy and standard MNI space, and automated removal of noise artefacts by independent components analysis (M. F. Glasser et al., 2013; Griffanti et al., 2014; Salimi-Khorshidi et al., 2014). We regressed the global signal and its temporal derivative from each rsfMRI time series and concatenated the four scans. We then computed the zero lag Pearson correlation between the concatenated time series from each pair of regions to derive the functional connectivity matrix, which we then Fisher’s z-transformed. We used the vectorised upper triangular of this functional connectivity matrix to predict cognition.

### Cognition

The NIH Toolbox Cognition Battery is an extensively validated battery of neuropsychological tasks (Carlozzi et al., 2017; Gershon et al., 2013; Heaton et al., 2014; Mungas et al., 2014; Tulsky et al., 2017; Weintraub et al., 2013; Weintraub et al., 2014; Zelazo et al., 2014) that assesses five cognitive domains: language, executive function, episodic memory, processing speed, and working memory through seven individual test instruments (Heaton et al., 2014). The specific tasks include Dimensional Change Card Sort Test, Flanker Inhibitory Control and Attention Test, Picture Sequence Memory Test, Picture Vocabulary Test, Oral Reading Recognition Test, List Sorting Working Memory Test, and Pattern Comparison Processing Speed (Heaton et al., 2014). Three composite scores are derived from participants’ scores on the NIH Toolbox Cognitive Battery tasks: Crystallised Cognition Composite, Fluid Cognition Composite, and Total Cognition Composite (Heaton et al., 2014). Crystallised cognition primarily represents language (vocabulary and reading decoding) abilities, while fluid cognition represents a wider range of higher-order cognitive processes including executive function (cognitive flexibility and inhibitory control and attention), episodic memory, working memory, and processing speed. These composite scores are based on initial factor analysis of the NIH Toolbox Cognition Battery. Specifically, the Crystallised Cognition Composite comprises the Picture Vocabulary and Oral Reading Recognition tests and assesses language and verbal skills. The Fluid Cognition Composite comprises scores on the Dimensional Change Card Sort, Flanker Inhibitory Control and Attention, Picture Sequence Memory, List Sorting Working Memory, and Pattern Comparison Processing Speed tests to broadly assess processing speed, memory, and executive functioning. The Total Cognition Composite combines the Crystallised and Fluid Cognition Composites. Composite scores tend to be more reliable/stable but do not capture variability in individual tasks (Heaton et al., 2014). In this study, we investigated the Crystallised, Fluid, and Total Cognition Composites, along with the individual scores from the seven tasks comprising them.

### Prediction of Cognitive Performance

We used functional connectivity to predict ten distinct outputs (three composite scores and seven task scores). For each prediction, we trained three distinct models: one sex-independent (trained on both male and female subjects), and two sex-specific (one trained on males, and one trained on females). For each model, we randomly shuffled and split the male and female subjects into train (80%) and test (20%) splits. We concatenated the male and female training sets for the sex-independent models, and kept them separate for the sex-specific models. We fit a linear ridge regression model on Scikit-learn (Pedregosa et al., 2011) using the training subset and tuned the regularisation parameter with five shuffled iterations of nested cross validation with three-fold inner and outer loops. We optimised the regularisation parameter in the inner loop and validated it in the outer loop. We took the median optimised hyperparameters from the five iterations to generate a single final model. We trained this model on the entire (sex-independent or sex-specific) training set, extracted feature weights, and evaluated the model’s prediction accuracy and explained variance on two distinct hold-out test sets: one test set comprised of male subjects and the other comprised of female subjects. Male and female train and test sets consisted of equal numbers of subjects. We quantify prediction accuracy as the Pearson correlation between the true and predicted values (J. W. Li et al., 2019). We repeated this using 100 unique train/test splits to generate a distribution of performance metrics.

### Model Significance

For each predictive model, we generated a corresponding null distribution to assess model significance as previously described (Dhamala et al., 2021; Parkes et al., 2021). We permuted the predicted variables (cognitive score) 25,000 times and then randomly split the data into train and test sets. For each of these 25,000 permutations, we trained and tested the model on the permuted data to obtain a null distribution of model performance. We assessed whether the original model’s performance was significantly non-zero by comparing the prediction accuracy from each of the original model’s 100 train/test splits to the median prediction accuracy from the null distribution. Specifically, the p-value for the model’s significance is the proportion of 100 original models that had prediction accuracies less than or equal to the median performance of the null model. We then corrected the p-values for multiple comparisons over all models (trained on both sexes, trained on males only, and trained on females only to predict ten distinct cognitive scores) and both test subsets (males only and females only) using the Benjamini-Hochberg False Discovery Rate (q=0.05) procedure (Benjamini & Hochberg, 1995).

### Model Comparisons

For each cognitive score, our workflow generated two distributions of 100 performance values: the first representing model performance when evaluated on only male individuals, and the second representing model performance when evaluated on only female individuals. For each cognitive score, we compared prediction performance across the male and female test sets using an exact test of differences (MacKinnon, 2009).

### Feature Importance

We adjusted feature weights to increase their interpretability as described in (Haufe et al., 2014). Briefly, for each iteration of a model, we used the feature weights, W, the covariance of the input variable (functional connectivity) in the training set, Σ_*x*_, and the covariance of the output variable (cognitive score) in the training set, Σ_*y*_, to extract the adjusted feature weights, A, as follows:

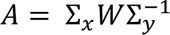

We then averaged the adjusted feature weights over the 100 iterations of each model to obtain feature importance matrices. Pairwise regional feature importances were mapped to the network level (Figure S1) by assigning each cortical region from the CoCo439 atlas to one of 17 networks from the Yeo 17-network parcellation (Yeo et al., 2011). Subcortical regions in the CoCo439 atlas were assigned to a subcortical network, and cerebellar regions to a cerebellar network. The average of the positive and negative feature importances of region pairs within and between the 17 networks were calculated separately; the result is a set of positive and negative importance of connections between and within the 17 networks. We evaluated the Pearson correlation between different models’ pairwise network-level feature importances, where positive and negative importances were considered together by concatenating them into a single vector. We also computed sex differences in positive and negative importance of connections between and within the 17 networks using an exact test of differences (MacKinnon, 2009).

## Results

An overview of our experimental workflow is shown in Figure 1. Please refer to the Methods section for details.

### Sex-Independent Models

Sex-independent models significantly predict Total and Crystallised Composite scores for both sexes, and Fluid Composite scores in males only, (corrected p<0.05). Within the crystallised domain, we significantly predict Picture Vocabulary scores in both sexes (corrected p<0.05), but only significantly predict Reading scores in females (corrected p<0.05). Within the fluid domain, we significantly predict Dimensional Change Card Sort, Picture Sequence Memory, and Processing Speed scores in males (corrected p<0.05), while we fail to significantly predict Flanker and List Sorting scores in males or females. Prediction accuracy for sex-independent models is shown in Figure 2 and Table 1. Explained variance for sex-independent models is shown in Figure S2 and Table S1.

**Figure 2:**
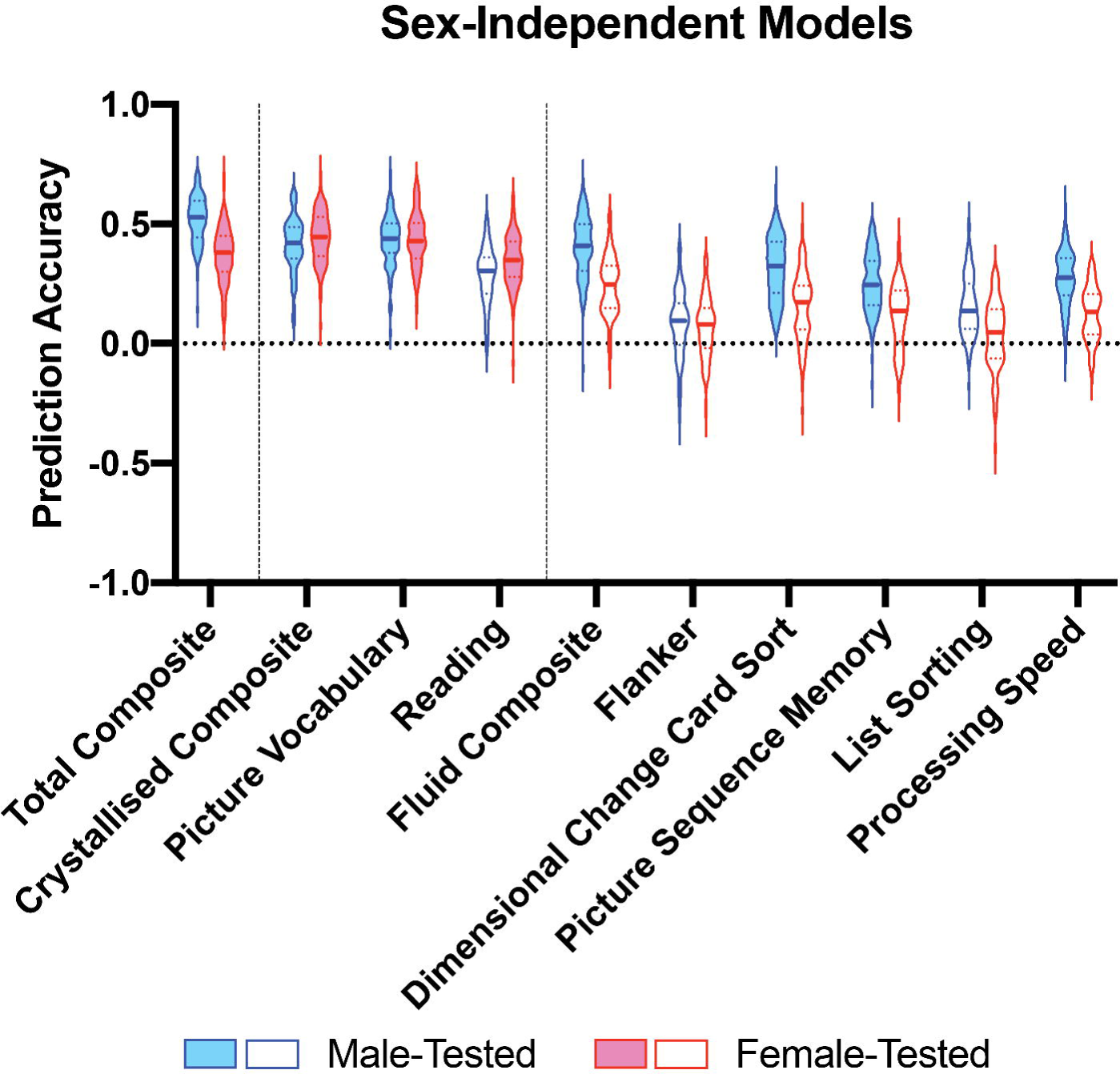
Violin plots of prediction accuracy (correlation between true and predicted cognitive scores) for sex-independent models predicting cognitive composite scores and individual task scores. Blue violins represent accuracy of models tested on male subjects and red represents of models tested on female subjects. The shape of the violin plots indicates the entire distribution of values, dashed lines indicate the median, and dotted lines indicate the interquartile range. Solid colour violin plots represent models that performed above chance levels based on permutation tests. Vertical dotted lines separate individual tests according to cognitive domain: general, crystallised, and fluid.

**Table 1:** Prediction accuracy (correlation between true and predicted cognitive scores) for sex-independent models predicting cognitive composite scores and individual task scores. Median prediction accuracy (interquartile range) is shown. Bolded prediction accuracy values denote that the model performed better than chance after corrections for multiple comparisons. * denotes p<0.05, ** denotes p<0.01, *** denotes p<0.001.

|  | Male-Tested | Female-Tested |
| --- | --- | --- |
| **Total Composite** | **0.53 (0.15) \*\*\*** | **0.38 (0.15) \*\*\*** |
| **Crystallised Composite** | **0.42 (0.13) \*\*\*** | **0.45 (0.16) \*\*\*** |
| **Picture Vocabulary** | **0.44 (0.12) \*\*\*** | **0.43 (0.15) \*\*\*** |
| **Reading** | 0.30 (0.15) | **0.35 (0.15) \*** |
| **Fluid Composite** | **0.41 (0.19) \*** | 0.25 (0.17) |
| **Flanker** | 0.10 (0.17) | 0.08 (0.16) |
| **Dimensional Change Card Sort** | **0.32 (0.21) \*\*\*** | 0.17 (0.18) |
| **Picture Sequence Memory** | **0.25 (0.18) \*** | 0.14 (0.20) |
| **List Sorting** | 0.14 (0.19) | 0.05 (0.20) |
| **Processing Speed** | **0.28 (0.15) \*** | 0.13 (0.17) |

### Sex-Specific Models

Sex-specific male-trained and female-trained models significantly predict Total Composite scores in both sexes (corrected p<0.05). Using female-trained models, we significantly predict Crystallised Composite scores in both sexes (corrected p<0.05), but fail to significantly predict Fluid Composite scores in either sex. Using male-trained models, we significantly predict Crystallised Composite scores in females and Fluid Composite scores in males (corrected p<0.05). Within the crystallised domain, we significantly predict Picture Vocabulary scores in both sexes using both sex-specific models (corrected p<0.05), but only significantly predict Reading scores in the opposite sex (corrected p<0.05). Within the fluid domain, we significantly predict Dimensional Change Card Sort in males using male-trained models (corrected p<0.05), but fail to significantly predict all Flanker, Picture Sequence Memory, List Sorting, and Processing Speed scores in either sex using either sex-specific model. Prediction accuracy for sex-specific models is shown in Figure 3 and Table 2. Explained variance for sex-specific models is shown in Figure S3 and Table S2.

**Figure 3:**
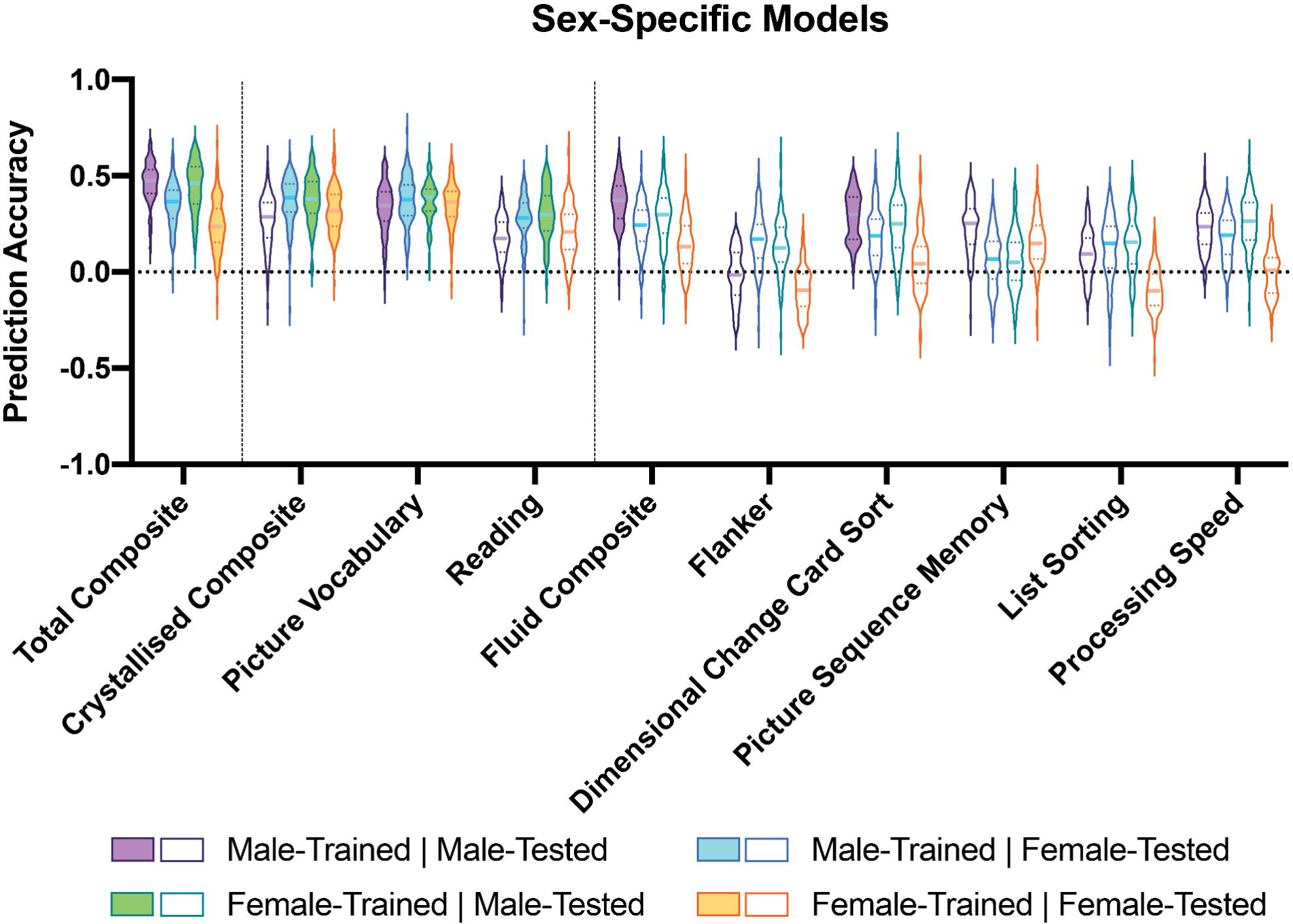
Violin plots of prediction accuracy (correlation between true and predicted cognitive scores) for sex-specific models predicting cognitive composite scores and individual task scores. Purple indicates results from models trained and tested on males; blue indicates results from models trained on males and tested on females; green indicates results from models trained on females and tested on males; and orange indicates results from models trained and tested on females. The shape of the violin plots indicates the entire distribution of values, dashed lines indicate the median, and dotted lines indicate the interquartile range. Solid colour violin plots indicate those models that performed above chance levels based on permutation tests. Vertical dotted lines separate individual tests according to cognitive domain: general, crystallised, and fluid.

**Table 2:** Prediction accuracy (correlation between true and predicted cognitive scores) for sex-specific models predicting cognitive composite scores and individual task scores. Median prediction accuracy (interquartile range) is shown. Bolded prediction accuracy values denote that the model performed better than chance after corrections for multiple comparisons. * denotes p<0.05, ** denotes p<0.01, *** denotes p<0.001.

|  | Male-Trained |  | Female-Trained |  |
| --- | --- | --- | --- | --- |
|  | Male-Tested | Female-Tested | Male-Tested | Female-Tested |
| **Total Composite** | **0.48 (0.12) \*\*\*** | **0.36 (0.14) \*** | **0.46 (0.19) \*\*\*** | **0.24 (0.17) \*** |
| **Crystallised Composite** | 0.29 (0.18) | **0.39 (0.14) \*** | **0.38 (0.16) \*\*\*** | **0.32 (0.16) \*** |
| **Picture Vocabulary** | **0.35 (0.15) \*** | **0.38 (0.16) \*\*\*** | **0.39 (0.11) \*\*\*** | **0.36 (0.13) \*** |
| **Reading** | 0.17 (0.16) | **0.28 (0.13) \*** | **0.30 (0.18) \*** | 0.21 (0.18) |
| **Fluid Composite** | **0.37 (0.17) \*** | 0.24 (0.16) | 0.30 (0.18) | 0.13 (0.19) |
| **Flanker** | -0.02 (0.22) | 0.17 (0.17) | 0.13 (0.17) | -0.10 (0.17) |
| **Dimensional Change Card Sort** | **0.30 (0.22) \*\*\*** | 0.19 (0.18) | 0.25 (0.22) | 0.04 (0.19) |
| **Picture Sequence Memory** | 0.25 (0.18) | 0.07 (0.19) | 0.05 (0.19) | 0.15 (0.17) |
| **List Sorting** | 0.09 (0.17) | 0.15 (0.21) | 0.15 (0.19) | -0.10 (0.16) |
| **Processing Speed** | 0.24 (0.16) | 0.19 (0.17) | 0.26 (0.19) | 0.01 (0.18) |

### Model Comparisons

Using exact tests of differences, we did not identify any significant differences in model performance between the sexes in the sex-independent models or the sex-specific models for any cognitive score.

### Feature Importance Comparisons

We correlated network-level feature importances between the sex-independent and sex-specific models (Figure 4). Feature importances between all pairs of sex-independent and sex-specific models are significantly correlated (corrected p<0.05). Features important in predicting the Total Composite, Crystallised Composite, and specific crystallised task scores from the sex-independent models are equally correlated to those from the male- and female-specific models. Features important in predicting the Fluid Composite and specific fluid scores are more strongly correlated to features important to predict those scores in males than in females. Features important in predicting each of the scores from the sex-specific models are generally more strongly correlated within sexes for different cognitive scores than across sexes for the same cognitive score; however, the correlations between models trained on different sexes is generally high. Features important in predicting the Total Composite score are correlated with features important to predict the Crystallised and Fluid composite scores and each of the individual task scores. Feature importance for predicting specific crystallised task scores are more strongly correlated with feature importance for predicting the Crystallised Composite score in females than they are in males. Features important for predicting specific fluid task scores are more strongly correlated to those important for predicting the Fluid Composite score in males than they are in females.

**Figure 4:**
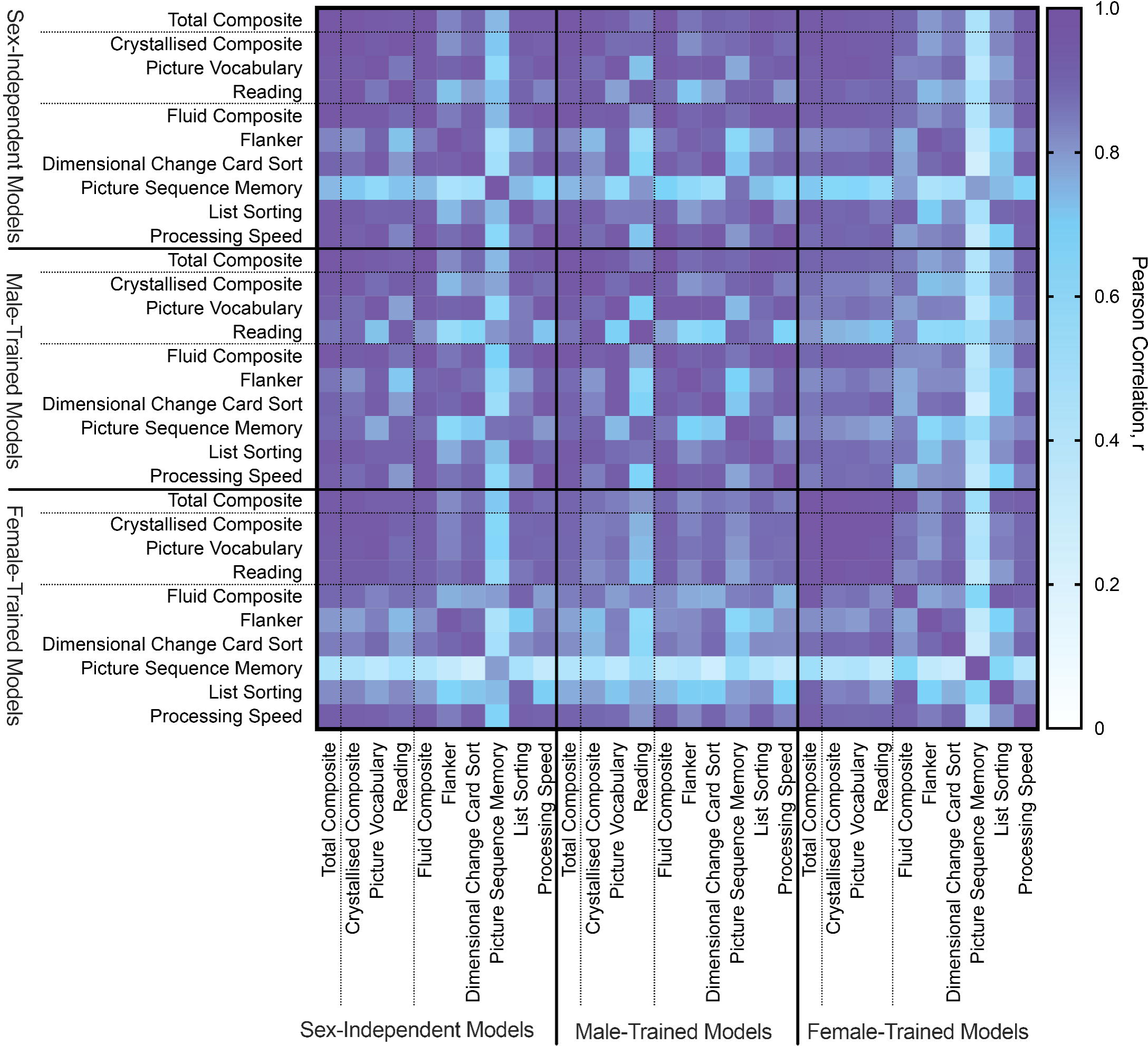
Pearson correlation of network-level feature importance for the sex-independent and sex-specific models predicting each cognitive score. Positive and negative network-level feature importance were computed by taking the positive and negative sums of the regional feature importance. Correlations were evaluated between the concatenated positive and negative network-level feature importances.

### Network-Level Feature Importance

Stronger functional connections between visual, dorsal attention, ventral attention, and temporal parietal networks are associated with higher crystallised abilities in males and females (Figure 5). Stronger functional connections within and between visual, dorsal attention, ventral attention, and temporal parietal networks, as well as within visual, dorsal attention, and default mode networks predict higher fluid abilities in females, while stronger functional connections between visual, ventral attention, and temporal parietal networks predict higher fluid abilities in males. Stronger functional connections within visual, somatomotor, and temporal parietal networks predict lower fluid and crystallized abilities in both sexes. Generally similar functional connections predict Picture Vocabulary and Reading scores in both sexes (Figure 6) as well as scores in individual fluid tasks, with the exception of List Sort and Picture Sequence scores (Figure S4). In females, stronger functional connections within visual, dorsal attention, control, and default mode networks predict higher List Sort scores, while stronger connections between those networks predict lower scores. In males, stronger connections between visual, dorsal attention, and ventral attention, as well as within dorsal attention, control, and default mode networks predict higher List Sort scores, while stronger connections within visual, somatomotor, and temporal parietal networks predict lower scores. Stronger functional connections within visual and temporal parietal networks predict higher Picture Sequence scores in females, while stronger connections within the default mode network, and between visual, dorsal attention, and ventral attention networks predict higher scores in males. Similar patterns of connectivity-cognition associations are also observed with the sex-independent models (Figure S5).

**Figure 5:**
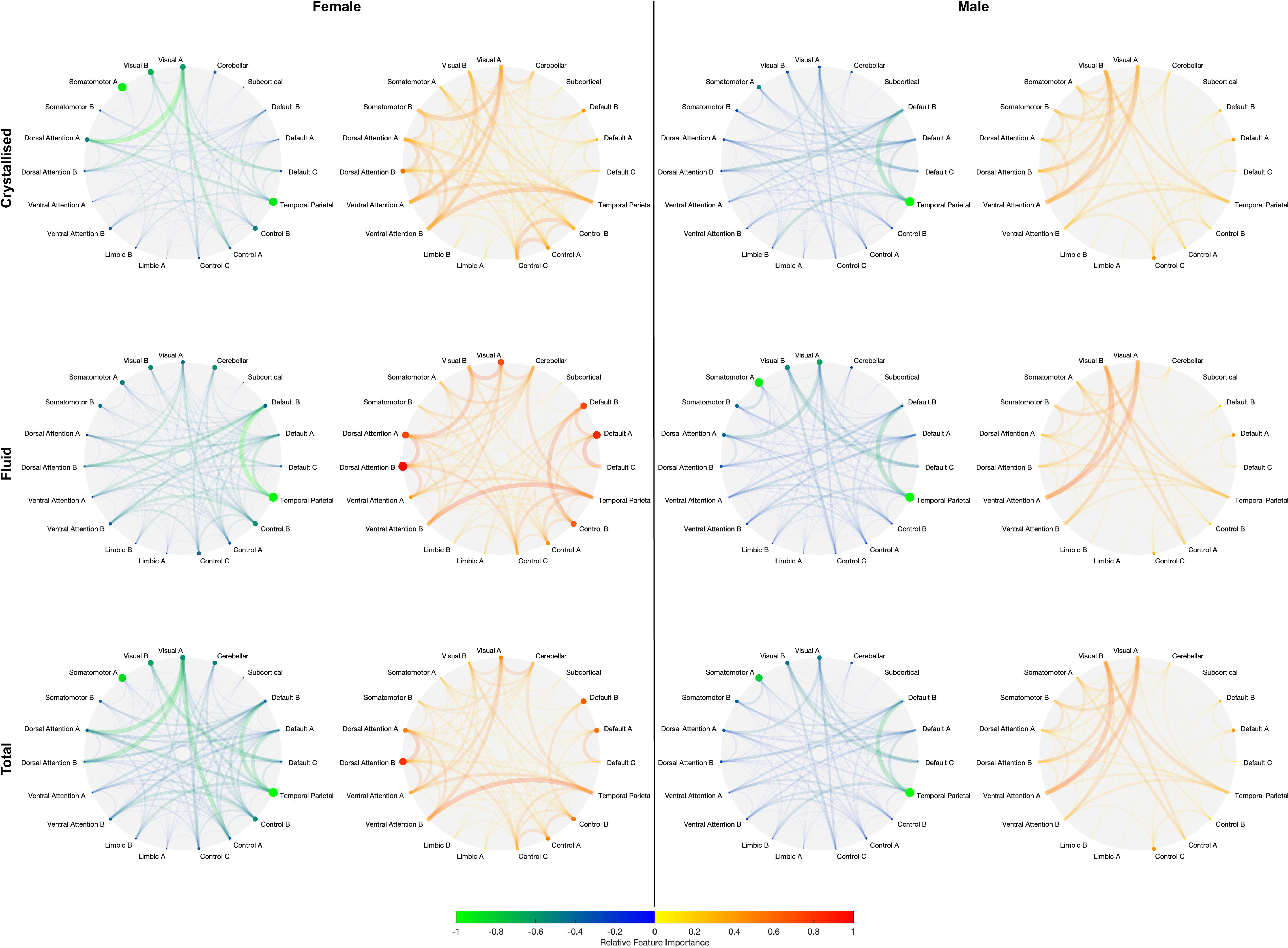
Network-level positive and negative feature importance for females (left two columns) and males (right two columns) to predict crystallised (top), fluid (middle), and total (bottom) cognition composites. Node radii and colour denote strength of intra-network feature importance. Edge thickness and colour denote strength of inter-network feature importance. Warmer colours are used for positive feature importance, and cooler colours for negative feature importance.

**Figure 6:**
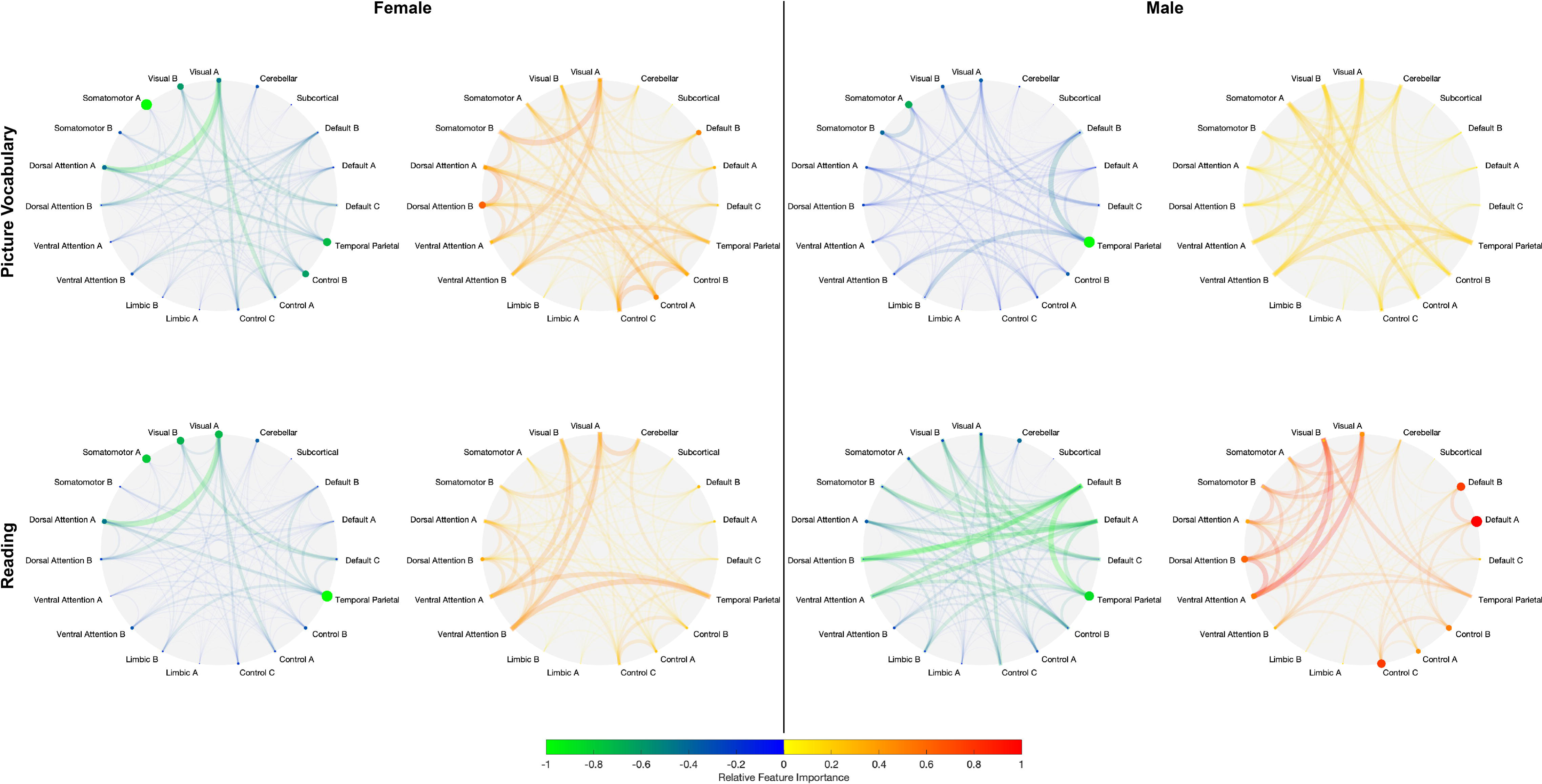
Network-level positive and negative feature importance for females (left two columns) and males (right two columns) to predict individual crystallised cognition task scores: picture vocabulary (top) and reading (bottom). Node radii and colour denote strength of intra-network feature importance. Edge weight and colour denote strength of inter-network feature importance. Warmer colours are used for positive feature importance, and cooler colours for negative feature importance.

### Sex Differences in Network-Level Feature Importance

Using exact tests of differences, we found there are no significant sex differences in the strength of the positive or negative associations between functional connectivity and the Crystallised Composite, Picture Vocabulary, Reading, or Flanker scores, but there are significant sex differences in the strength of the positive and negative associations between functional connectivity and Total Composite, Fluid Composite, Card Sort, List Sort, Picture Sequence, and Processing Speed scores (Figure S6). Specifically, females generally exhibit stronger negative connectivity-cognition associations (i.e., stronger functional connections = lower cognitive scores), while males generally exhibit stronger positive connectivity-cognition associations (i.e., stronger functional connections = higher cognitive scores).

## Discussion

In this study, we quantified sex-independent and sex-specific relationships between functional connectivity and cognition. Using whole brain resting-state functional connectivity, we predicted individual crystallised and fluid abilities in 392 healthy young adults. First, we find sex-independent models predict with equivalent accuracy crystallised abilities in both sexes but predict fluid abilities more accurately in males. Second, we show sex-specific models perform comparably when predicting crystallised abilities within and between sexes, but generally fail to predict fluid abilities in either sex, except for the Fluid Composite and Dimensional Change Card Sort score in males. Third, we demonstrate that sex-specific models predicting crystallised and fluid abilities generally rely on shared functional connections within and between distinct cortical networks. Together, our findings largely suggest that shared neurobiological features predict general and specific crystallised abilities in both sexes.

Crystallised cognition represents language abilities, while fluid cognition represents executive function, memory, and processing speed. Prior work has shown Total and Crystallised Composite scores are more predictable than the Fluid Composite (Dhamala et al., 2021) but that work did not investigate whether the same is true for specific tasks within the cognitive domains or whether these results hold equally among males and females. In this current work, we replicate and expand upon those previous findings.

Results from our sex-independent models suggest they might be capturing shared relationships between functional connectivity and crystallised abilities in males and females, but male-specific relationships between functional connectivity and fluid abilities. This is supported by our observation that connectivity-cognition relationships for fluid abilities from the sex-independent models more closely resemble those from the male-specific models than the female-specific models. Results from our sex-specific models provide additional support for our findings from the sex-independent models, as we find that connectivity-cognition relationships for crystallised abilities and overall cognition are generally shared between the sexes. We also observe an even greater inability to predict fluid abilities with our sex-specific models compared to our sex-independent models, which could be in part due to the decreased sample size in the sex-specific models. The general lack of predictability observed for fluid abilities in both types of models may be underscored by individual differences in the signal-to-noise ratio of the specific brain-behaviour relationships. Fluid abilities are more susceptible to factors including sleep, stress, and mood which directly influence executive functions and memory and less stable within an individual over time (Nilsson et al., 2005; O’Neill, Kamper-DeMarco, Chen, & Orom, 2020; Salthouse, 2010). This contradicts prior reports of successful prediction of fluid intelligence from functional connectivity (Finn et al., 2015). However, it is noting that even though our models do not perform better than chance (as evaluated by comparing to a null distribution), our Fluid Composite prediction accuracies are generally comparable to those previously reported as significant (evaluated using Pearson’s correlation) (Finn et al., 2015). Other prior work has also demonstrated that fluid intelligence, as well as other behavioural variables, can be successfully predicted using white matter functional connectivity at accuracies comparable to those reported in this study (J. Li, Biswal, et al., 2020; J. Li, Chen, et al., 2020). Our smaller sample size and choice of significance evaluation method may explain our inability to successfully predict fluid intelligence and the differences in the conclusions we draw from our results. Moreover, despite initially matching participants across sex by cognitive scores, we also observe a greater inability to predict cognitive abilities in females compared to males, and this may be due to sex differences in the variances of the cognitive scores. Of the ten cognitive measures predicted in this study, the Fluid Cognition Composite, Flanker, and Processing Speed scores have significantly different variances across the sexes. Specifically, male scores for those cognitive measures had significantly larger variances than female scores (corrected p <0.05; data not shown). Using the sex-independent models, the Fluid Cognition Composite and the Processing Speed scores, were predicted above chance levels in males but not in females, while the Flanker predictions were comparable to chance levels for both sexes. A lower variance in the scores within the females means a more restricted range of scores making it less likely that a significant association can be identified. Additionally, it may result in a lower signal-to-noise ratio in females, thus make the scores harder to predict. Similar results demonstrating sex differences in predictions of fluid intelligence have previously been published (Greene, Gao, Scheinost, & Constable, 2018). Specifically, they showed that models using resting-state functional connectivity to predict fluid intelligence in children/adolescents and adults tend to perform better in males than in females across different edge thresholds (Greene et al., 2018). Moreover, they also showed that predicting fluid intelligence using emotion task-based models significantly outperformed working memory task-based models in females, while working memory task-based models significantly outperformed emotion task-based models in males, and suggested that there exist fundamental sex differences in the spatial distribution and modulation of networks related to fluid intelligence (Greene et al., 2018).

Our understanding of cognitive sex differences and brain-behaviour relationships have widely shifted over the decades. While research has confirmed some differences, many others have been refuted (Halpern, 2013; Miller & Halpern, 2014). Two similar studies to date investigating sex-specific brain-behaviour relationships have reported contradictory findings. Implementing a connectome-based prediction modelling approach, Jiang et al observed no differences in prediction accuracy between males and females when predicting IQ using functional connectivity (Jiang, Calhoun, Cui, et al., 2020). In a second study from the group, they demonstrated IQ was more predictable in females than in males (Jiang, Calhoun, Fan, et al., 2020). In this current work, our sex-independent models comparably predict overall cognition and crystallised abilities in males and females, but better predict some fluid abilities in males compared to females while failing to predict other fluid tasks in either sex altogether. In this study, we implemented a nested cross validation approach with 100 different randomised splits of the data to generate a distribution of performance accuracy measures. Previous studies relied on integrating feature selection with a leave-one-out cross validation approach resulting in a single accuracy value for the model and distinct features being used to predict the output variable for each subject (Jiang, Calhoun, Cui, et al., 2020; Jiang, Calhoun, Fan, et al., 2020). Due to these methodological differences, our prediction accuracy results cannot be directly compared to prior work. However, it is worth noting that our sex-specific models comparably predict overall and crystallised aspects of cognition in males and females, supporting one of the previous studies (Jiang, Calhoun, Cui, et al., 2020) but contradicting the other (Jiang, Calhoun, Fan, et al., 2020).

In this study, we find connections within and between distinct cortical networks are crucial to predict cognition, and these features are shared between the sexes, contradicting extant literature implementing sex-specific models (Jiang, Calhoun, Cui, et al., 2020; Jiang, Calhoun, Fan, et al., 2020). More specifically, we find stronger connections between the visual, dorsal attention, ventral attention, and temporal parietal networks predict higher crystallised and fluid ability scores in both sexes, while stronger connections within visual, somatomotor, and temporal parietal networks predict lower crystallised and fluid ability scores in both sexes. While some differences in male and female models’ feature importances exist, their correlations are moderate to high (R = 0.6-0.9). Specifically, we observe that there exist significant sex differences in the strength of the positive and negative associations between functional connectivity and Total Composite, Fluid Composite, Card Sort, List Sort, Picture Sequence, and Processing Speed scores. While males and females share positive (i.e., stronger functional connections predict higher cognitive scores) and negative (i.e., stronger functional connections predict weaker cognitive scores) connectivity-cognition associations, females exhibit stronger negative relationships between distributed network connectivity patterns and the cognitive scores while males exhibit stronger positive relationships. Hence, while males and females share the same connectivity-cognition relationships, the strength of those relationships may vary between the sexes. However, the List Sort, Picture Sequence, and Processing Speed models performed worse than chance for predictions in both sexes, and the Fluid Composite and Card Sort models only performed better than chance in males, limiting the relevance of this finding. We also demonstrate that feature importance correlations, within and between sexes, are stronger for tasks within the crystallised domain than tasks within the fluid domain or tasks between the two domains. This is likely related to the models’ overall lower accuracies in predicting fluid abilities; if the models are not reliably mapping functional connectivity to fluid abilities, there will be more noise in their feature importance, resulting in lower correlations across models. Our results contradict findings from prior work identifying distinct correlates of cognition in males and females. In one study, authors reported the top 100 functional connections to predict IQ in males and females are distinct with only three overlapping features (Jiang, Calhoun, Fan, et al., 2020). In a second study, authors found male IQ was more strongly correlated with functional connectivity in left parahippocampus and default mode network, while female IQ was more strongly correlated with functional connectivity in putamen and cerebellar network (Jiang, Calhoun, Cui, et al., 2020). This discrepancy in findings could be due to model differences, particularly in the cross-validation, feature selection, and inference choices, or the choice of cognitive score.

## Limitations

In this study, we trained and tested sex-independent and sex-specific models on 196 male and 196 female subjects, all unrelated. Over each of the 100 unique train/test splits, we ensured the same set of male/female subjects were in the training and testing subsets for the sex-independent and the male/female-specific models. Maintaining this consistency of subjects allowed us to maintain the variance within the subjects, but also resulted in our sex-independent models being trained and tested on twice as many subjects as our sex-specific models. Prior work has demonstrated that fluid abilities are more difficult to predict than crystallised abilities (Dhamala et al., 2021). In this study, we found sex-independent models were able to predict some fluid abilities above chance levels in males, but sex-specific models generally did not perform above chance levels for either sex. The inherent difficulty in predicting fluid abilities, combined with the lower number of subjects for the sex-specific models, may explain why many of our sex-specific models performed poorly. In this study, our main goal was to evaluate whether the models differed in their predictions of cognitive abilities between males and females rather than between the models themselves. However, future work in this area should explore whether sex-independent and sex-specific models differ from one another when training sample sizes are consistent.

Many researchers studying cognitive differences between males and females compare group averages between the sexes. While this approach can yield insightful results pertaining to general sex differences, their relevance to individual cognitive abilities in males and females is limited. Genetic, hormonal, cultural, and psychosocial factors can influence sex-related and sex-independent individual differences in functional connectivity and cognition (Cosgrove, Mazure, & Staley, 2007; Miller & Halpern, 2014). Here, we sought to uncover whether relationships between functional connectivity and cognition are shared between the sexes or are distinct. Our results largely suggest shared network connectivity features equally predict cognitive abilities in males and females. However, we must acknowledge that here, due to the limitations of the data set, we can only consider individuals’ sex but not their gender identity or fluidity. Our society projects distinct gender roles onto males and females paving the way for a lifetime of gender-differentiated experiences (Eliot, 2011). These distinct social factors may drive gender differences in brain-behaviour relationships, even in the absence of sex differences, that our study is not designed to capture. Future work in this area should aim to collect and integrate data about gender identity and fluidity so we can better understand how relationships between connectivity and cognition may or may not vary with gender.

Many machine learning models based on neuroimaging data struggle with generalisability due to differences in study sites, scanner types, and scan parameters. The models we have designed in this study were only trained, validated, and tested on data from the Human Connectome Project. Although we implement a nested cross validation approach and evaluate our models with 100 distinct train/test splits, the results we report may not be entirely comparable or generalisable to other datasets. Future studies should aim to integrate data from multiple sites to address this limitation.

## Conclusion

A comprehensive understanding of neurobiological markers that underlie cognitive abilities within and across sexes is necessary if we are to understand sex-specific effects of aging and illness on cognition. Here, we implement predictive modelling approaches to explore sex-independent and sex-specific relationships between functional connectivity and cognitive abilities. We report three main findings. We demonstrate that sex-independent models comparably capture relationships between connectivity and crystallised abilities in males and females, but only successfully capture relationships between connectivity and fluid abilities in males. We find sex-specific models comparably predict crystallised abilities within and between sexes, but fail to predict fluid abilities in either sex. Finally, we find that stronger connections between visual, dorsal attention, ventral attention, and temporal parietal networks predict higher crystallised and fluid ability scores, and stronger connections within visual, somatomotor, and temporal parietal networks predict lower crystallised and fluid ability scores in both sexes. Taken together, this suggests that brain-behaviour relationships are shared between the sexes and rely on overlapping network connectivity within and between cortical structures.

## Acknowledgements

This work was supported by the following National Institutes of Health (NIH) grants: R21 NS104634-01 (AK) and R01 NS102646-01A1 (AK). The sponsor did not have any role in the study design, the analysis, or interpretation of the data; the writing of the report; or the decision to submit the manuscript for publication. Data were provided and made available by the Human Connectome Project, WU-Minn Consortium (Principal Investigators: David Van Essen and Kamil Ugurbil; 1U54MH091657) funded by the 16 NIH Institutes and Centres that support the NIH Blueprint for Neuroscience Research; and by the McDonnell Centre for Systems Neuroscience at Washington University.

## Citation Gender Diversity Statement

Recent work in neuroscience and other fields has identified a bias in citation practices such that papers from women and other minorities are under-cited relative to the number of such papers in the field (Caplar, Tacchella, & Birrer, 2017; Chakravartty, Kuo, Grubbs, & McIlwain, 2018; Dion, Sumner, & Mitchell, 2018; Dworkin et al., 2020; Maliniak, Powers, & Walter, 2013; Thiem, Sealey, Ferrer, Trott, & Kennison, 2018). Here we sought to proactively consider choosing references that reflect the diversity of the field in thought, form of contribution, gender, and other factors. We used classification of gender based on the first names of the first and last authors (Dworkin et al., 2020), with possible combinations including male/male, male/female, female/male, and female/female. Excluding self-citations to the first and last authors of our current paper, the references contain 45.2% male/male, 16.7% male/female, 21.4% female/male, and 16.7% female/female. We look forward to future work that could help us to better understand how to support equitable practices in science.

## Author Contributions

Conceptualization, E.D., A.K.; Methodology, E.D., K.W.J., A.K.; Software, E.D.; Investigation, E.D., A.J., A.K.; Formal Analysis, E.D., A.K.; Resources, A.K; Data Curation, E.D., K.W.J.; Writing – Original Draft, E.D.; Writing – Review & Editing, E.D., K.W.J., A.J., A.K.; Visualisation, E.D., K.W.J.; Supervision, A.K.; Funding Acquisition, A.K.

## Data Availability Statement

The data used are openly available as part of the Human Connectome Project at https://www.humanconnectome.org/study/hcp-young-adult/document/1200-subjects-data-release.

