## Supplementary Materials for "Shared functional connections within and between cortical networks predict cognitive abilities in adult males and females"

### Figure S1: Mapping from CoCo 439 atlas to Yeo network.

Mapping from the CoCo 439 parcellation to the Yeo 17-network parcellation with added networks for subcortical and cerebellar regions.

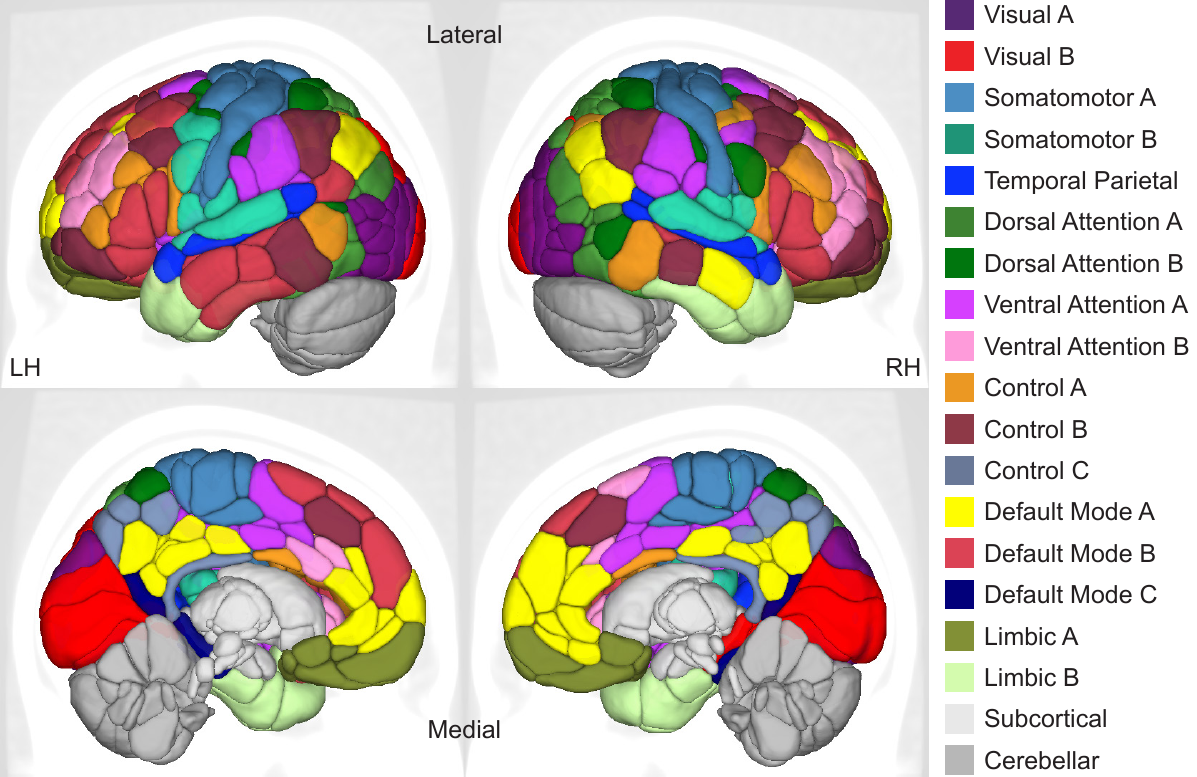

Figure S2: Violin plots of explained variance for sex-independent models predicting cognitive composite scores and individual task scores. Blue violins represent accuracy of models tested on male subjects and red represents of models tested on female subjects. The shape of the violin plots indicates the entire distribution of values, dashed lines indicate the median, and dotted lines indicate the interquartile range. Solid colour violin plots represent models that performed above chance levels based on permutation tests. Vertical dotted lines separate individual tests according to cognitive domain: general, crystallised, and fluid.

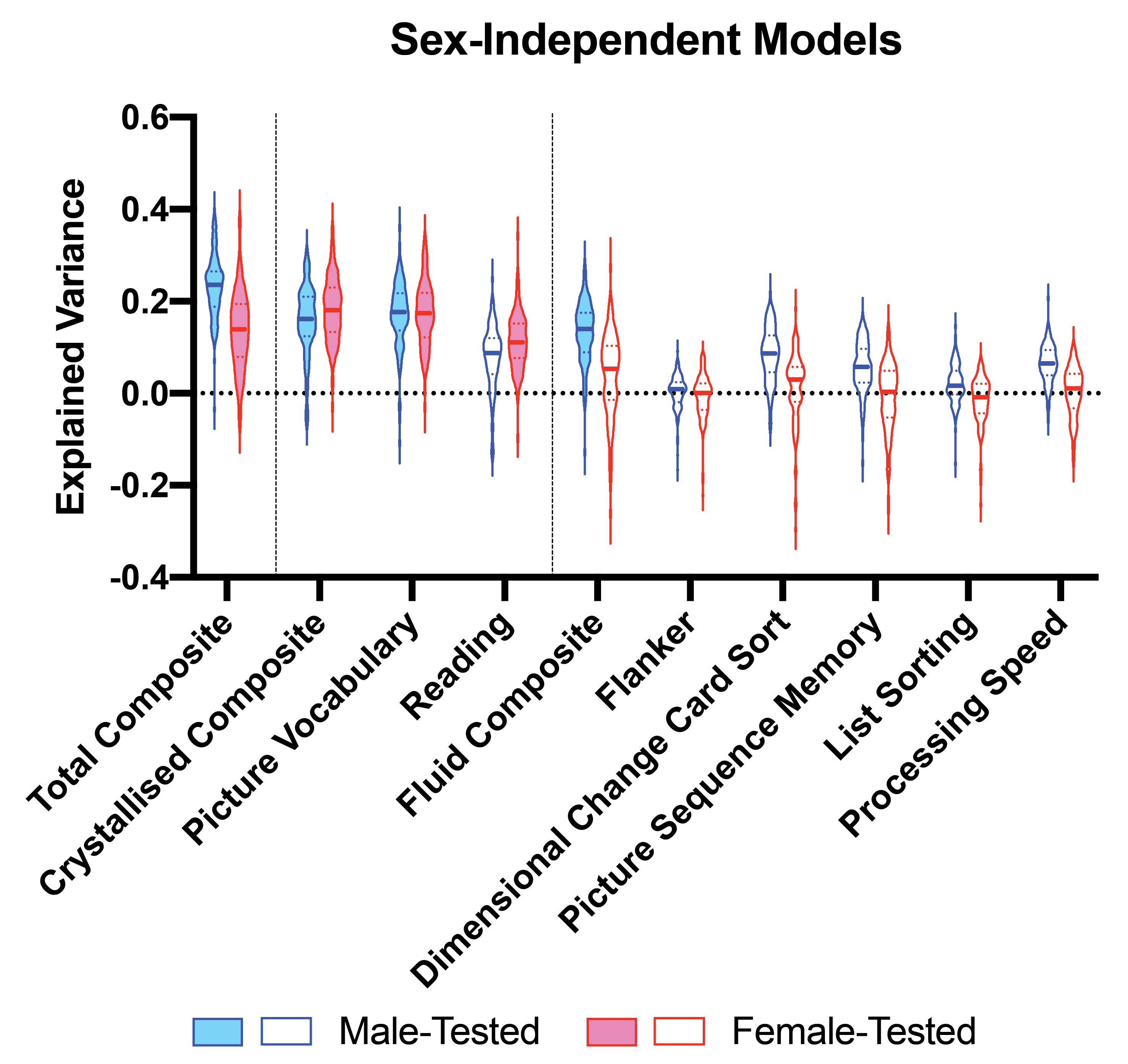

Table S1: Explained variance for sex-independent models predicting cognitive composite scores and individual task scores. Median explained variance (%) (interquartile range) is shown. Bolded explained variance values denote that the model performed better than chance after corrections for multiple comparisons. * denotes p<0.05, ** denotes p<0.01, *** denotes p<0.001.

|  | Male-Tested | Female-Tested |
| --- | --- | --- |
| **Total Composite** | **23.5 (7.7) ***** | **13.9 (11.0) *** |
| **Crystallised Composite** | **16.1 (8.3) *** | **18.0 (9.5) ***** |
| **Picture Vocabulary** | **17.6 (7.7) *** | **17.3 (9.0) ***** |
| **Reading** | 8.7 (7.9) | **11.1 (7.4) *** |
| **Fluid Composite** | **14.0 (8.3) *** | 5.3 (11.5) |
| **Flanker** | 0.9 (4.0) | 0.1 (5.3) |
| **Dimensional Change Card Sort** | 8.6 (7.9) | 2.9 (7.5) |
| **Picture Sequence Memory** | 5.7 (7.3) | 0.3 (9.6) |
| **List Sorting** | 1.6 (4.9) | -0.9 (6.4) |
| **Processing Speed** | 6.4 (5.4) | 1.0 (7.3) |

Figure S3: Violin plots of explained variance for sex-specific models predicting cognitive composite scores and individual task scores. Purple indicates results from models trained and tested on males; blue indicates results from models trained on males and tested on females; green indicates results from models trained on females and tested on males; and orange indicates results from models trained and tested on females. The shape of the violin plots indicates the entire distribution of values, dashed lines indicate the median, and dotted lines indicate the interquartile range. Solid colour violin plots indicate those models that performed above chance levels based on permutation tests. Vertical dotted lines separate individual tests according to cognitive domain: general, crystallised, and fluid.

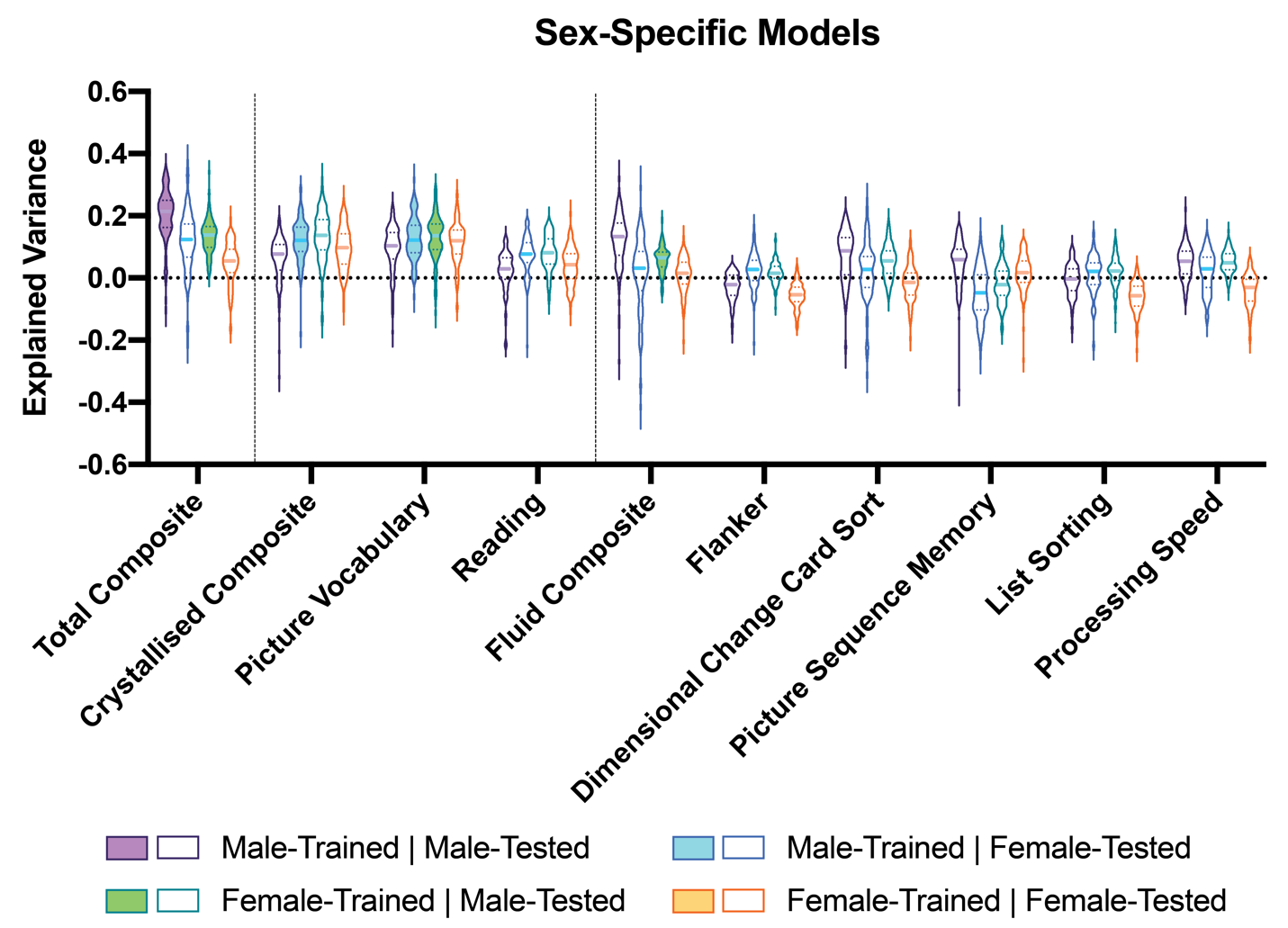

Table S2: Explained variance for sex-specific models predicting cognitive composite scores and individual task scores. Median explained variance (%) (interquartile range) is shown. Bolded explained variance values denote that the model performed better than chance after corrections for multiple comparisons. * denotes p<0.05, ** denotes p<0.01, *** denotes p<0.001.

|  | Male-Trained | | Female-Trained | |
| --- | --- | --- | --- | --- |
|  | Male-  Tested | Female-  Tested | Male-  Tested | Female-  Tested |
| **Total Composite** | **21.4 (8.8) *** | 12.4 (10.6) | **13.8 (6.5) ***** | 5.5 (7.4) |
| **Crystallised Composite** | 7.8 (8.1) | **12.1 (7.8) *** | 13.8 (9.5) | 9.7 (9.7) |
| **Picture Vocabulary** | 10.4 (8.2) | **12.2 (8.7) ***** | **13.6 (8.1) *** | 12.0 (7.7) |
| **Reading** | 3.0 (6.8) | 7.7 (6.2) | 8.1 (7.9) | 4.3 (7.8) |
| **Fluid Composite** | 13.3 (10.4) | 3.1 (12.2) | **6.5 (4.7) *** | 1.5 (6.9) |
| **Flanker** | -2.2 (6.3) | 2.7 (6.4) | 1.5 (3.9) | 5.3 (4.6) |
| **Dimensional Change Card Sort** | 8.7 (11.8) | 2.8 (10.0) | 5.5 (7.1) | -1.4 (6.9) |
| **Picture Sequence Memory** | 5.9 (8.7) | -4.8 (11.1) | -2.1 (7.5) | 1.7 (6.7) |
| **List Sorting** | -0.3 (7.0) | 2.1 (6.8) | 2.2 (5.5) | -5.7 (6.3) |
| **Processing Speed** | 5.4 (7.2) | 2.9 (9.5) | 4.9 (5.0) | -3.0 (7.0) |

Figure S4: Network-level positive and negative feature importance for females (left two columns) and males (right two columns) to predict individual fluid cognition task scores. Node radii and colour denote strength of intra-network positive and negative feature importance. Edge weight and colour denote strength of inter-network positive and negative feature importance. Warmer colours are used for positive feature importance, and cooler colours for negative feature importance.

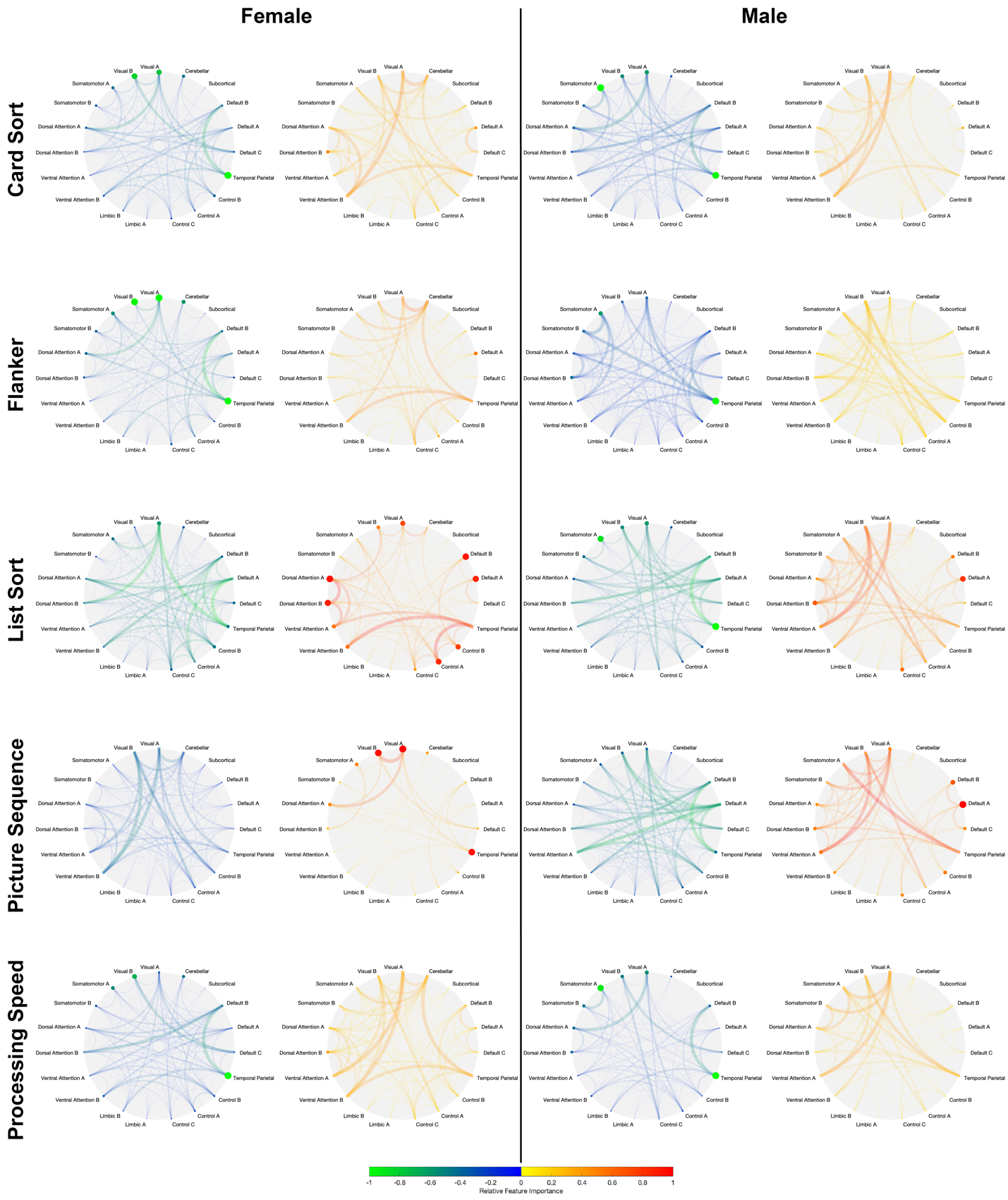

Figure S5: Network-level positive and negative feature importance from sex-independent models to predict cognitive composite and individual cognition task scores. Node radii and colour denote strength of intra-network positive and negative feature importance. Edge weight and colour denote strength of inter-network positive and negative feature importance. Warmer colours are used for positive feature importance, and cooler colours for negative feature importance.

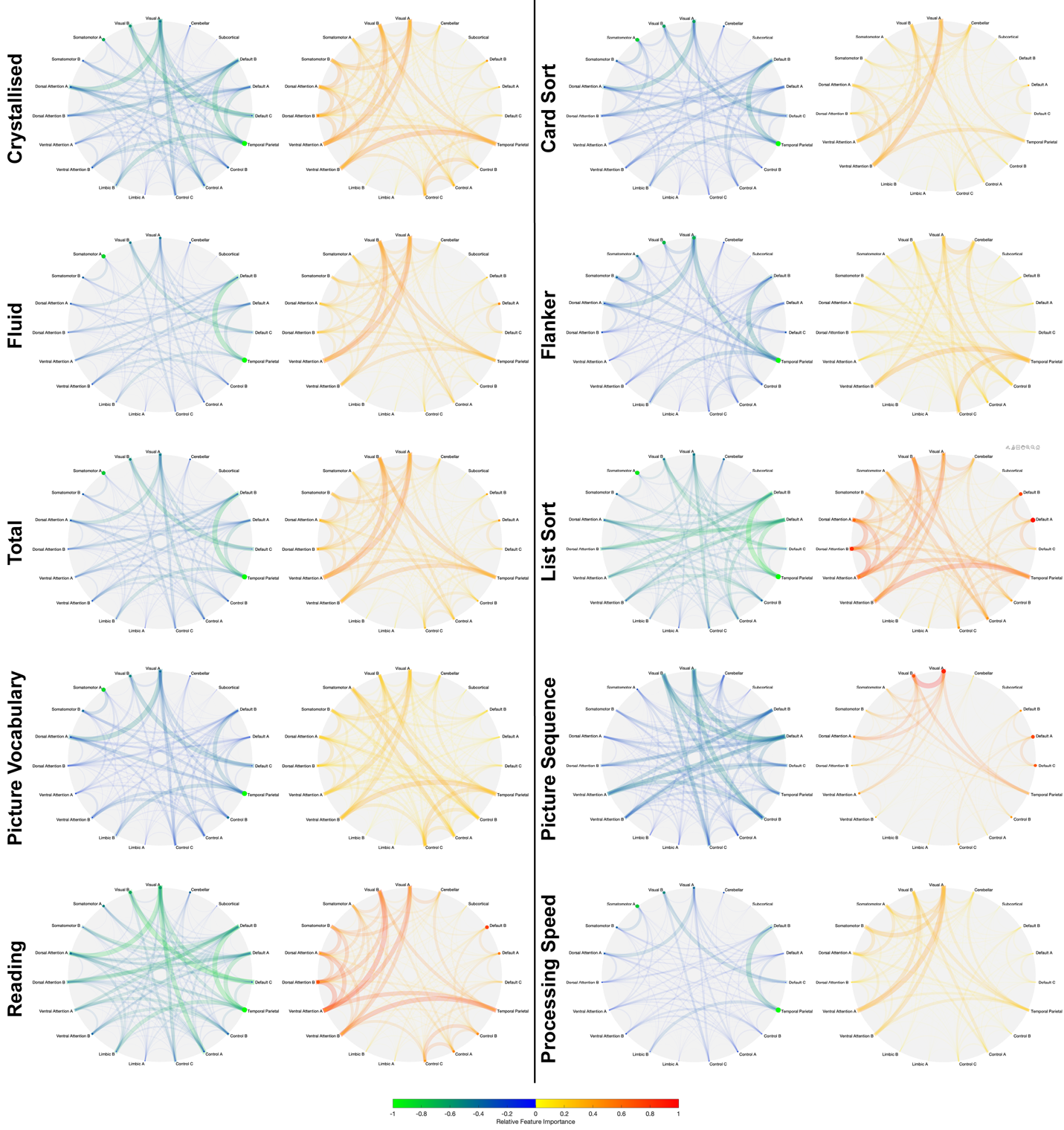

Figure S6: Network-level difference in means in positive and negative feature importance between male- and female- specific models to predict cognitive composite and individual cognition task scores. Node radii/edge weight and colour denote relative strength of difference in the positive and negative feature importances. Warmer colours denote that female-specific models have a higher feature importance (i.e., more positive, or less negative) than male-specific models and cooler colours denote that male-specific models have a higher feature importance than female-specific models. Only edges that are significantly different between the sexes (computed using an exact test for differences and corrected for multiple comparisons) are shown. Plots are not shown for Crystallised Composite, Picture Vocabulary, Reading, or Flanker scores because no significant differences were observed in the positive or negative feature importances for those predictions.

­

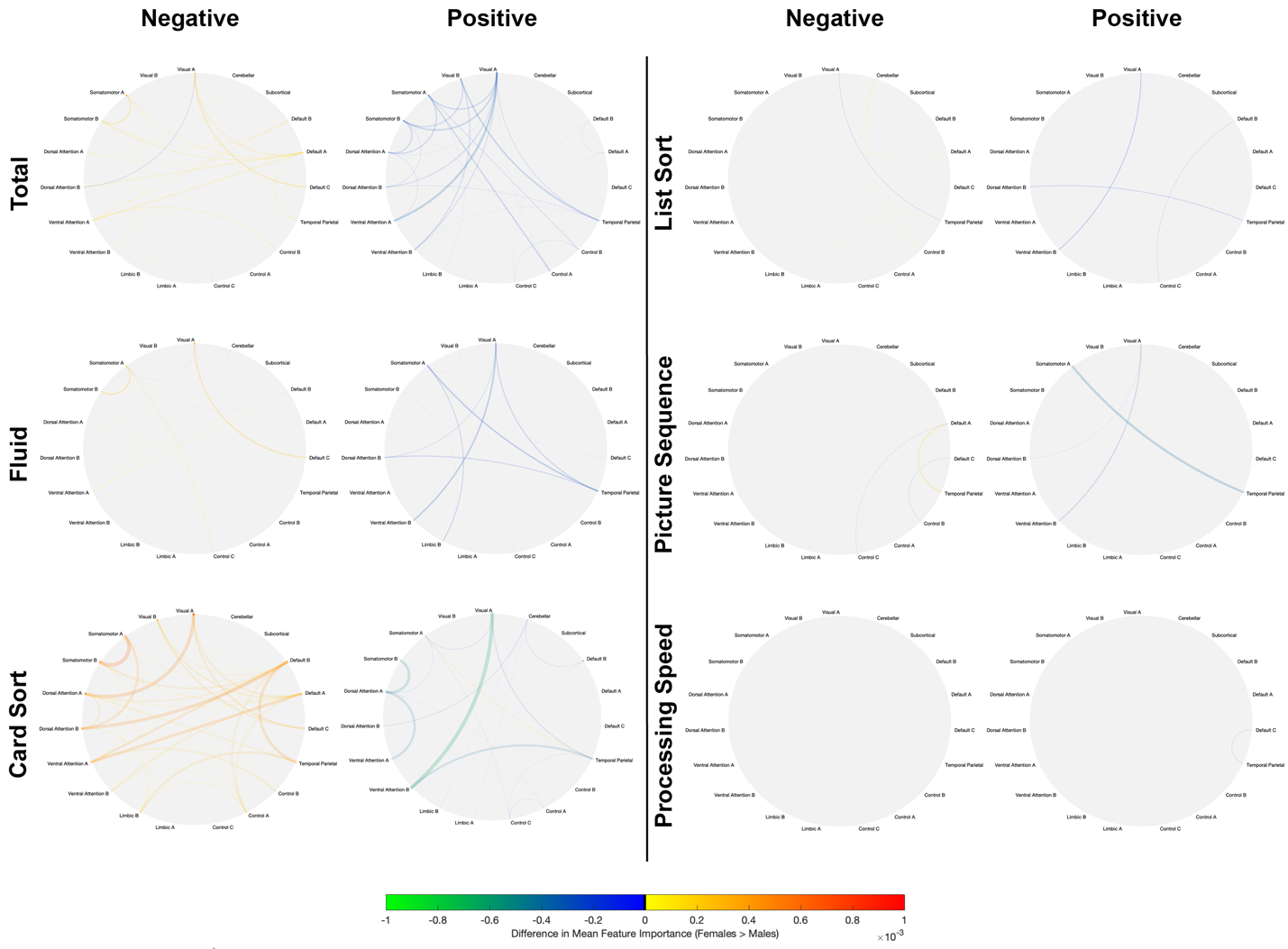
